# Relationships between balance performance and connectivity of motor cortex with primary somatosensory cortex and cerebellum in middle aged and older adults

**DOI:** 10.1101/2024.03.29.587335

**Authors:** Ashwini Sansare, Thamires N.C. Magalhaes, Jessica A. Bernard

## Abstract

Connectivity of somatosensory cortex (S1) and cerebellum with the motor cortex (M1) is critical for balance control. While both S1-M1 and cerebellar-M1 connections are affected with aging, the implications of altered connectivity for balance control are not known. We investigated the relationship between S1-M1 and cerebellar-M1 connectivity and standing balance in middle-aged and older adults. Our secondary objective was to investigate how cognition affected the relationship between connectivity and balance. Our results show that greater S1-M1 and cerebellar-M1 connectivity was related to greater postural sway during standing. This may be indicative of an increase in functional recruitment of additional brain networks to maintain upright balance despite differences in network connectivity. Also, cognition moderated the relationship between S1-M1 connectivity and balance, such that those with lower cognition had a stronger relationship between connectivity and balance performance. It may be that individuals with poor cognition need increased recruitment of brain regions (compensation for cognitive declines) and in turn, higher wiring costs, which would be associated with increased functional connectivity.

## 1. Introduction

About 30-40% of the community-dwelling older adults (OA) over the age of 65 fall once a year (Shumway-Cook and Woollacott, 1985; Rubenstein, 2006), and these rates nearly double in individuals over the age of 75 (Rubenstein, 2006). Falls not only lead to severe injuries, hospitalizations, and limited independence, but also lead to psychological consequences such as fear of falling, anxiety, and depression (Bates et al., 1995; Thomas et al., 2008; Painter et al., 2012). Understanding how aging affects the brain regions and networks that underlie balance control is a critical first step in developing targeted interventions for balance rehabilitation in OA.

Control of balance requires contributions from both motor and sensory systems as well as constant online control of the motion of the body. While prior research on the aging brain and balance has focused mainly on the motor aspect i.e., the primary motor cortex (M1) and supplementary motor area (Wu et al., 2007; Langan et al., 2010; Taubert et al., 2016), two critical brain regions that communicate with M1 for sensorimotor control of balance are somatosensory area (S1) and cerebellum. The synergistic activity between M1 and S1 is important for sensorimotor control; M1 interacts with S1 to integrate sensory feedback for movements (Salimi et al., 1999; Gardner et al., 2007) whereas S1 receives input from M1 to implement an online processing of somatosensory signals during movements (Umeda et al., 2019). With advanced age, the interactions between S1 and M1 are affected. The resting state functional connectivity between S1 and M1 is lower in OA compared to young adults (Cheng and Lin, 2013; He et al., 2017; Cassady et al., 2019). Moreover, the decline in S1-M1 connectivity is associated with poor upper limb motor function, such as tapping and gripping, in OA (Seidler et al., 2015a). While these studies have probed the link between S1-M1 connectivity and upper limb function in OA, the implications of reduced S1-M1 connectivity for balance control in advanced age are not known.

With respect to the cerebellum and sensorimotor control, the cerebellum implements an internal model that predicts sensory consequences of motor actions and refines the motor output when there are errors in these sensory predictions (Wolpert et al., 1998; Bastian, 2006). It plays a crucial role in controlling, initiating, and adjusting upright balance in humans (Thach and Bastian, 2004). The cerebellum interacts with the M1 through unique closed-loop circuits via the thalamus (Kelly and Strick, 2003; Salmi et al., 2010; Bernard et al., 2012, 2016). This cerebellar-M1 connectivity is essential for motor control and learning in the lower (Jayaram et al., 2011) and upper limbs (Schlerf et al., 2012; Spampinato et al., 2017). With aging, there is a reduction in cerebellar volume (Bernard & Seidler, 2013) as well as differences in cerebello-cortical network connectivity. Specifically, OA show widespread degradation in the cerebellar-M1 connectivity compared to young adults (Bernard et al., 2013; Rurak et al., 2022). Moreover, greater cerebellar-M1 connectivity correlated with the greater upper limb dexterity and higher confidence in balance control (Bernard et al., 2013), and faster walking performance (Rurak et al., 2022). As is the case with S1-M1 connectivity, there is no research probing the direct link between age-related declines in cerebellar-M1 connectivity and postural sway during standing.

Compared to young adults, OA demonstrate greater postural sway area in standing, which implies poor balance control (Baloh et al., 1994; Muir et al., 2013), and is an indicator of fall risk (Baczkowicz et al., 2008; Howcroft et al., 2017). Further, postural sway in OA increases with the removal of visual input i.e., eyes closed or by decreasing the somatosensory input from feet i.e., standing on foam surface, or a combination of both i.e., eyes closed on foam surface (Lord et al., 1991; Anson et al., 2019; Fujio and Takeuchi, 2021). Prior work on neuroimaging and balance has shown that poorer balance performance is correlated with brain structure changes such as greater declines in white matter (Starr et al., 2003; Sullivan et al., 2009; Van Impe et al., 2012) and reduced gray matter volume in superior parietal cortex and cerebellum (Rosano et al., 2007) in OA. Cerebello-cortical resting state connectivity has been associated with postural control in youth at clinical high risk for psychosis (a sample that also shows increased postural sway area) (Bernard et al., 2014), though critically this was an adolescent and young adult sample. Work investigating such connectivity in advanced age is lacking. Indeed, none of these studies investigated age-related differences in brain connectivity between the areas that contribute to sensorimotor control and its effect on balance. Hence, it is yet to be determined how age differences in S1-M1 and cerebellar-M1 connectivity are related to standing postural sway in advanced age.

Lastly, there are complex interactions between cognition and balance control during aging. Poor cognitive function is associated with worse balance performance potentially due to reduced attentional resources (Bahureksa et al., 2016; Li et al., 2018). Specifically, as OA multi-task with the addition of cognitive or sensory challenges, there is competition for limited neural resources, leading to trade-offs in either balance and walking performance or cognitive performance(Szturm et al., 2013). However, there is no research that probes how cognition may moderate the relationship between the age-related differences in network connectivity with M1 and balance.

The purpose of this study was to investigate the relationship between balance control and resting state S1-M1 and cerebellar-M1 connectivity in middle-aged and older adults. We hypothesized that reduced connectivity would be related to greater postural sway, which is indicative of poorer balance. As a secondary objective, we also investigated whether cognition moderated the relationship between S1-M1 and cerebellar-M1 connectivity and balance. We hypothesized that poor cognition would be related to a stronger relationship between connectivity and balance. Finally, to explore which connections of S1 and cerebellum with brain regions other than M1 are associated with balance control, we performed exploratory whole brain analyses by investigating the relationship between balance performance and the functional connectivity of S1 and cerebellum with whole brain.

## 2. Methods

### 2.1. Study sample

A total of 138 participants were recruited for the study (mean age ± SD = 58 ± 13, 71 females). We recruited participants from three age groups; early middle age (early MA): 35 to 50 years, n=47, mean age ± SD = 42 ± 5, 23 females; late middle-age (late MA): 51 to 64 years, n=51, mean age ± SD = 58 ± 5, 32 females; and older adults (OA): 65 years and above, n= 40, mean age ± SD = 72 ± 6, 19 females. Each participant underwent a comprehensive assessment, where their balance was assessed under four conditions (see section 2.2 Balance task for details) and cognitive function was assessed through Montreal Cognitive Assessment (MOCA) (Nasreddine et al., 2005). Following the behavioral visit, participants returned for a magnetic resonance imaging (MRI) session. Exclusion criteria included a history of neurological disease, stroke, or a formal diagnosis of psychiatric illness (e.g., depression or anxiety), as well as contraindications for the brain imaging environment. All study procedures received approval from the Institutional Review Board at Texas A&M University, and written informed consent was obtained from each participant before initiating any data collection. These data were collected as part of a larger longitudinal study on cerebellar aging, and only baseline data from those that completed both the imaging and behavioral visit are included here.

### 2.2. Balance task

Postural sway was assessed using an Advanced Mechanical Technology Incorporated Accusway force platform (Watertown, MA). Participants were asked to stand as still as possible with their hands crossed across their chest. Vision (eyes open/closed) and foot position (feet apart/closed) were varied such that participants completed the task under four conditions, eyes open with open base i.e. with feet shoulder width apart (EOOB), eyes closed with open base (ECOB), eyes open with close base i.e. with feet close to each other (EOCB) and eyes closed with closed base i.e. with feet close to each other (ECCB). Participants completed one trial of each condition. The center of pressure (COP) was recorded for two minutes with a sample rate of 50 Hz. To isolate the low-frequency postural sway, we applied a 9th order Butterworth filter with a 20 Hz cutoff frequency. The 95% confidence interval of COP area were calculated using previously established methods to quantify postural sway (Oliveira et al., 1996).

### 2.3. Imaging acquisition

Participants underwent structural and resting-state MRI using a Siemens Magnetom Verio 3.0 Tesla scanner equipped with a 32-channel head coil. For structural MRI, we obtained a high-resolution T1-weighted 3D magnetization prepared rapid gradient multi-echo (MPRAGE) scan with the following specifications: repetition time (TR) of 2400 ms, acquisition time of 7 minutes, and voxel size of 0.8 mm³. Additionally, a high-resolution T2-weighted scan was acquired with a TR of 3200 ms, acquisition time of 5.5 minutes, and voxel size of 0.8 mm³, both employing a multiband acceleration factor of 2. For resting-state imaging, we conducted four blood-oxygen-level-dependent (BOLD) functional connectivity (fMRI) scans, each lasting 6 minutes, resulting in a total of 24 minutes of resting-state imaging. The fMRI scans employed a multiband factor of 8, 488 volumes, a TR of 720 ms, and voxel dimensions of 2.5 mm³.

The scans were acquired with alternating phase encoding directions, encompassing two anterior-to-posterior scans and two posterior-to-anterior scans. Throughout the fMRI scans, participants were instructed to lie still with their eyes open, fixating on a central cross. The overall image acquisition process took approximately 45 minutes, including a 1.5-minute localizer. The scanning protocols were adapted from multiband sequences developed by the Human Connectome Project (HCP) (Harms et al., 2018) and the Center for Magnetic Resonance Research at the University of Minnesota, ensuring compatibility for future data sharing and enhancing reproducibility.

### 2.4. Functional connectivity analysis

The images underwent several preprocessing steps to prepare them for further analysis. First, they were converted from DICOM to NIFTI format and organized according to the Brain Imaging Data Structure (BIDS, version 1.6.0) using the bidskit docker container (version 2021.6.14, https://github.com/jmtyszka/bidskit). Next, a single volume was extracted from two oppositely coded BOLD images to estimate B0 field maps using the split tool from the FMRIB Software Library (FSL) package (Jenkinson et al., 2012). Subsequently, the anatomical and functional images were preprocessed using fMRIPrep (version 20.2.3; for detailed methods see https://fmriprep.org/), which involves automated procedures to align the functional volume with the anatomical image, correct for motion, correct field map distortions, segment the anatomical image into distinct tissues (e.g., gray matter, white matter, cerebrospinal fluid), remove the skull from the anatomical image, normalize the data to a common space, align motion-corrected functional volumes with the normalized anatomical image and apply spatial smoothing.

After preprocessing with fMRIPrep, we continued the rest of the analyses using the CONN toolbox (version 21a) (Whitfield-Gabrieli & Nieto-Castanon, 2012). This includes processing to remove noise and artifacts and improve the quality of the data. Denoising in CONN typically involves several steps, including the removal of motion signals and regression of confounding signals (such as signals from ventricles, white matter, and global signals). Motion information from fMRIPrep was available in CONN for this process. A 0.008-0.099 Hz bandpass filter was applied to remove high-frequency noise. The denoising step helps improve the quality of functional connectivity data by reducing artifacts and increasing sensitivity to detect genuine functional connectivity patterns in the brain.

In our first-level analyses, we included cerebellum regions of interest (ROIs) spherical seeds: Lobules V, VI, VII, VIII, and X (all right hemisphere) (more details in *ROI selection* section). Additionally, we included two customized cortical seeds—one in the S1 and one in the M1 areas of the cerebral cortex, responsible for sensory processing (S1) and voluntary movement control (M1) (see *ROI selection* section)—to explore cerebello-cortical connectivity. The S1 and M1 seeds were made in the lower limb area using the homunculus because we expect the lower limb to contribute to the standing balance control rather than the upper limb, face, or other regions of the body. Finally, we extracted first-level ROI-to-ROI correlation results to examine connectivity between the lobules and S1, as well as between the lobules and M1.

#### 2.4.1. ROI selection

We utilized the MRIcron Brodmann area atlas (www.nitrc.org/projects/mricron) as a reference for our custom ROIs (S1 and M1). Originally developed by Damasio and Damasio (1989) (Damasio & Damasio, 1989) for lesion mapping, this atlas has been updated with more recent findings from the Van Essen laboratory (Van Essen et al., 2000). The updated atlas includes maps of the Brodmann areas along with their corresponding Talairach coordinates. Considering the sensory homunculus, a topographic representation of the body’s sensory distribution in the cerebral cortex, we defined coordinates in area 4 (part of the precentral gyrus, representing M1-leg location) and areas 3, 1, and 2 (parts of the postcentral gyrus, representing S1-leg location). These coordinates and the spherical seeds created are detailed in Figure 1 and Table 1.

**Figure 1.**
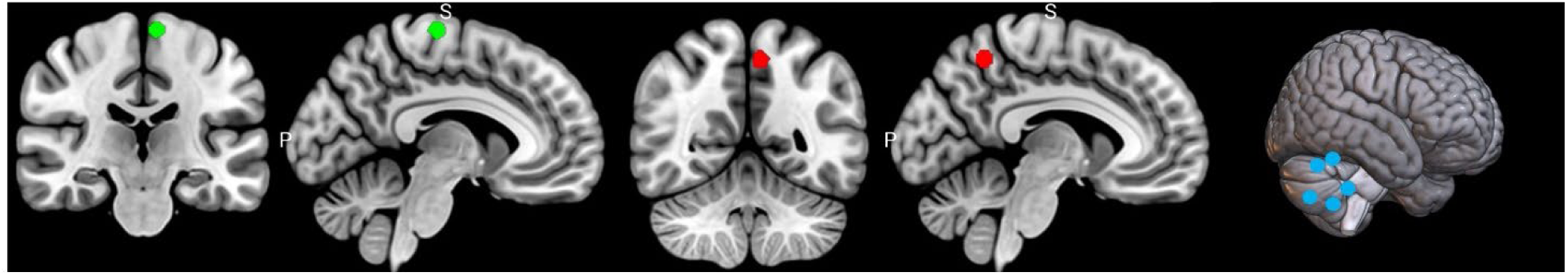
All the regions of interest (ROIs), with the leg area for M1 located at the bottom and highlighted in green, while the leg area for S1 is highlighted in red. Additionally, the seeds in the cerebellum region of the right hemisphere are represented in blue.

**Table 1:** MNI coordinates for the M1, S1 and cerebellar seeds used in analyses.

| ROIs | X | Y | Y |
| --- | --- | --- | --- |
| Left Cerebral Cortex |  |  |  |
| Left M1 | -7.91 | -21.52 | 70.56 |
| Left S1 | -7.17 | -53.97 | 53.6 |
| Right Cerebellar Lobules |  |  |  |
| V | 15 | -52 | -18 |
| VI | 25.3 | -59.9 | -25.3 |
| VII | 28.2 | -65.8 | -51.1 |
| VIII | 18 | -51 | -54 |
| X | 22.3 | -37.8 | -44.5 |

Spherical seeds were generated for cerebellar subregions (Lobules V, VI, VII, VIII, and X) using the SUIT atlas coordinates as a reference (Diedrichsen et al., 2009), with localization to the right hemisphere. The seeds, each specified to have a diameter of 5mm, were created using the Marsbar program (www.marsbar-toolbox.github.io), a MATLAB (www.mathworks.com) toolbox.

### 2.5. Statistical Analysis

To test how aging affected the balance performance on the four different balance conditions, we first performed a repeated measures ANOVA using balance condition (EOOB, ECOB, EOCB, ECCB) as a within subject factor and age (early MA, late MA and OA) as a between group factor. Pairwise post hoc comparisons were performed using Bonferroni corrections. Next, to determine the relationships between brain connectivity and balance, we performed bivariate correlations in SPSS (Released 2023. IBM SPSS Statistics for Windows, Version 29, Armonk, NY: IBM Corp) between connectivity (S1-M1, Lobule V-M1, Lobule VI-M1, Lobule VII-M1, Lobule VIII-M1 and Lobule X-M1) and balance performance on the ECCB condition. To limit multiple comparisons, we chose *a priori* to focus our analyses in this paper on the bivariate correlations with just one condition that would present the greatest challenge to postural stability. Bivariate correlations between connectivity and the remaining three balance conditions (EOOB, EOCB, and ECOB) were not statistically significant and are reported in Supplementary Table 1 for completeness. To determine whether one predictor or a combination of two or more predictors were best at predicting the balance performance on ECCB, data were further analyzed using a direct-entry multiple regression analysis (MRA), using all our connectivity measures as predictors and ECCB as the outcome.

Next, to investigate the role of cognition in the relationship between connectivity and balance, we divided our sample into two groups, high cognition, with MOCA scores in the normal range of 26-30 (n = 97) and low cognition, with MOCA scores between 20-25 (n = 41). We then performed bivariate correlations separately for high and low cognition groups for only those connectivity measures that significantly predicted ECCB performance in our MRA. A moderation analysis was then also conducted using PROCESS MACRO (Hayes, 2017), an extension of SPSS, with balance as the dependent variable, connectivity as the predictor, and cognition (ordinal variable: high and low cognition) as the moderator. Further, because declines in cognition can be in part due to age, which might make it difficult to parse out whether it is truly cognition moderating this relationship or the effects of age on cognition that are indirectly affecting the relationship between connectivity and balance, we ran a moderated moderation model that investigated whether age (Z) was moderating how cognition (W) affected the relationship between connectivity (X) and balance (Y) (Figure 2).

**Figure 2:**
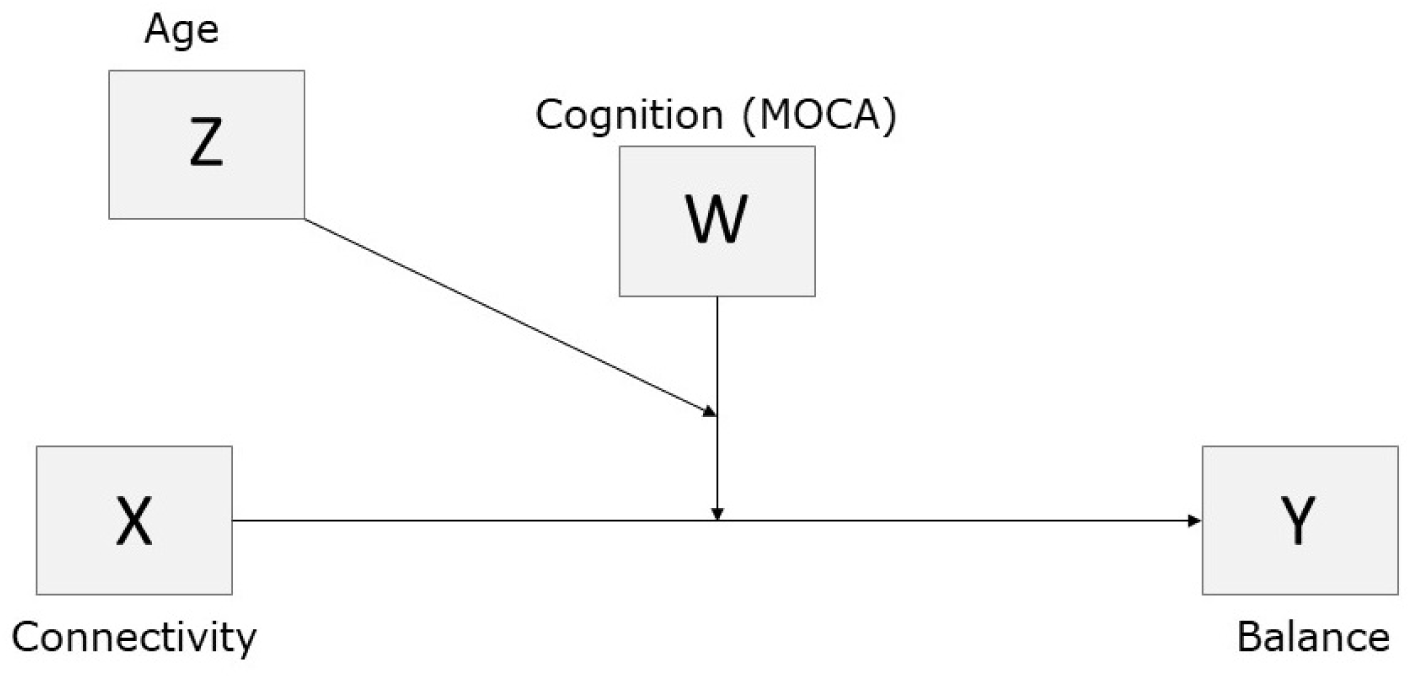
Representative model for the moderated moderation analysis with connectivity (X) as the independent variable, 95% COP area during balance task as the dependent variable (Y), cognition (high/low MOCA scores) as the moderator (W) and age (Z) as the second moderator.

Lastly, we conducted exploratory analyses of the relationship between balance performance in each of the four conditions and S1 and cerebellar connections with the whole brain by calculating seed-to-voxel resting state functional connectivity using the CONN toolbox. All analyses were first thresholded at p<0.001 at voxel level, followed by a cluster-level FDR-corrected p<0.01. The more stringent FDR-corrected p<0.01 at the cluster level was used to account for the additional comparisons in this exploratory analysis. For anatomical labeling of the MNI coordinates outputted by CONN, we used the label4MRI R package (https://github.com/yunshiuan/label4MRI) which uses the AAL (automatic anatomical labeling) atlas (Rolls et al., 2020).

## 3. Results

The descriptive statistics for age, MOCA scores, connectivity and balance measures are displayed in Table 2. While our initial sample had an n=138, we have missing data for some participants who either missed the visit for balance data collection or MRI session, or due to technical issues (lack of taring or “zeroing” the force plate before the participant stepped on the force plate) leading to incorrect balance data that had to be excluded from our analyses. This resulted in a different sample size for each balance condition as shown in the table below.

**Table 2:** Mean and Standard Deviations for all independent and dependent variables.

|  | n | Mean | Std Dev |
| --- | --- | --- | --- |
| S1-M1 | 138 | -0.0371 | 0.1032 |
| Lob V-M1 | 138 | -0.0081 | 0.08156 |
| Lob VI-M1 | 138 | -0.0249 | 0.08783 |
| Lob VII-M1 | 138 | -0.0147 | 0.09967 |
| Lob VIII-M1 | 138 | -0.0495 | 0.10154 |
| Lob X-M1 | 138 | -0.1059 | 0.12885 |
| EOOB ($m^2$) | 133 | 0.00018 | 0.00019 |
| ECOB ($m^2$) | 136 | 0.0002 | 0.00017 |
| EOCB ($m^2$) | 131 | 0.00043 | 0.00024 |
| ECCB ($m^2$) | 134 | 0.00058 | 0.00033 |
| MOCA score | 138 | 26.7319 | 2.13944 |
| Age (years) | 138 | 56.81 | 13.311 |

### 3.1. Effect of aging on balance

Our results from the ANOVA showed that there was a significant main effect for balance condition (F= 91.742, p < 0.001, ***η***^2^ = 0.689) but not for age (p = 0.182), nor an age group by condition interaction (p = 0.706), indicating that irrespective of the age, participants from all three groups had significantly different performance on the four balance conditions (Figure 3).

**Figure 3:**
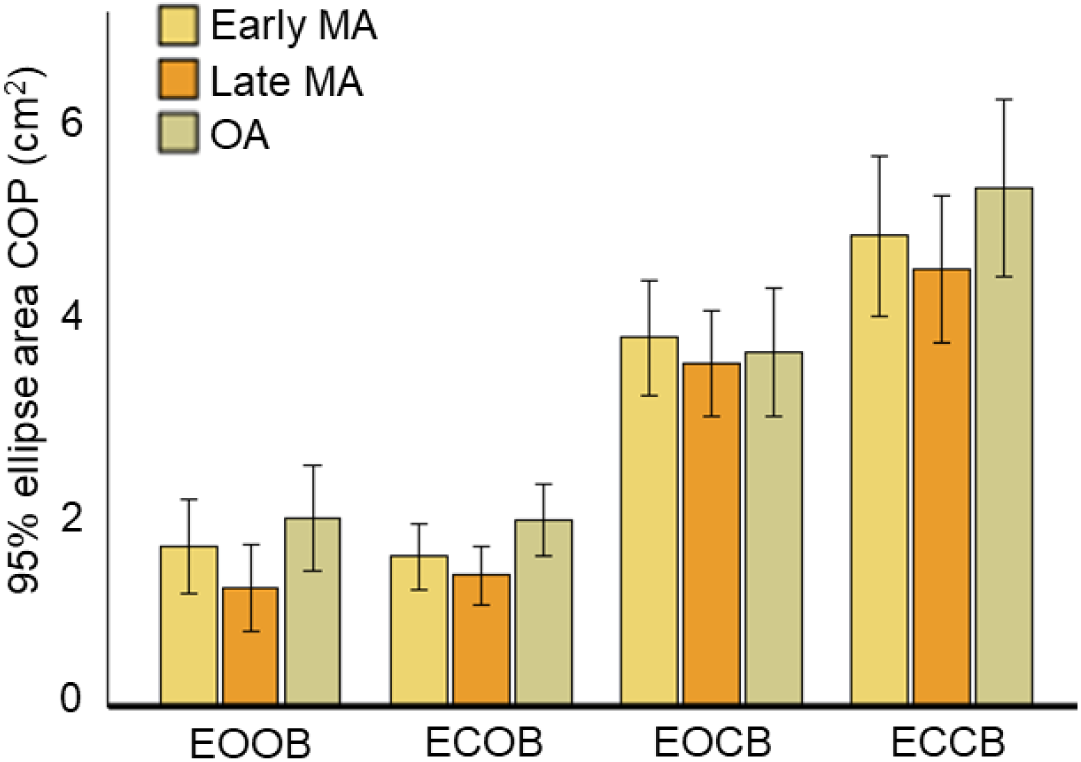
Postural sway depicted by the 95% ellipse area for center of pressure (COP) for the four balance conditions, eyes open (EO), eyes closed (EC), eyes open with closed balance i.e. narrow base of support (EOCB), eyes closed with closed balance (ECCB). Error bars indicate 95% confidence interval.

Post hoc tests revealed that the ECCB had significantly higher postural sway compared to the remaining three conditions (p < 0.001), followed by EOCB which had a significantly higher postural sway (p < 0.001) compared to the conditions with feet apart (EOOB, ECOB) and a significantly lower sway (p < 0.001) compared to the ECCB condition (Supplementary Table 2).

### 3.2. Relationship between connectivity and balance

Table 3 shows the p-values and correlation coefficients (r) for the relationships between connectivity and balance. Connectivity between S1-M1 and Lobule VII-M1 showed significant correlations with balance performance.

**Table 3.** Bivariate correlations between connectivity and balance performance on the most difficult condition (ECCB).

|  | S1-M1 | Lob V-M1 | Lob VI-M1 | Lob VII-M1 | Lob VIII-M1 | Lob X-M1 |
| --- | --- | --- | --- | --- | --- | --- |
| Pearson Correlation | .255** | -0.106 | -0.046 | .238** | 0.031 | -0.055 |
| Sig. (2-tailed) | 0.003 | 0.225 | 0.6 | 0.006 | 0.724 | 0.531 |

\* Correlation is significant at the 0.05 level (2-tailed).
\*\* Correlation is significant at the 0.01 level (2-tailed).

Results from the multiple linear regression are summarized in Table 4, with the overall model being statistically significant (F (6, 126) = 2.521, p = 0.024). S1-M1 connectivity (p = 0.026) and Lobule VIIa-M1 connectivity (p = 0.028) made statistically significant unique contributions to predicting performance on the ECCB condition.

**Table 4:** Regression analysis summary for connectivity variables predicting balance performance.

|  | S1-M1 | Lob V-M1 | Lob VI-M1 | Lob VII-M1 | Lob VIII-M1 | Lob X-M1 |
| --- | --- | --- | --- | --- | --- | --- |
| $\beta$ | 0.197 | -0.078 | -0.010 | 0.199 | -0.020 | -0.048 |
| t | 2.246 | -0.877 | -0.108 | 2.230 | -0.215 | -0.537 |
| p value | 0.026* | 0.382 | 0.914 | 0.028* | 0.830 | 0.592 |
| $sr^2$ | 0.051 | 0.002 | < 0.001 | 0.025 | < 0.001 | 0.004 |
| $f^2$ | 0.040 | 0.006 | < 0.001 | 0.040 | < 0.001 | 0.002 |

Note: $R^2 = .327$ ( $N = 133$ , $p = .010$ ), $\beta$ = standardized co-efficient, $sr^2$ = squared semi-partial coefficient, $f^2$ = Cohen's (1988) effect size statistic for multiple regression analyses. \* $p < .05$

Results (using the squared semi-partial coefficients) showed the S1-M1 connectivity made the largest unique contribution i.e., the contribution of the predictor after the effects of all other predictors are controlled. Lastly, effect sizes calculated for the predictors using Cohen’s f^2^ (Cohen, 2013), where values of .02 represent a small effect, values of .15 equal a medium effect, and values of .35 denote a large effect, show that S1-M1 and LobVIIa-M1 connectivity each had a small-to-medium effect size (respectively, f^2^ = .040 and 0.040) in predicting balance performance.

### 3.3. Effects of cognition on the relationship between connectivity and balance

Next, to investigate how cognition might affect the relationship between connectivity and balance, we performed bivariate correlations between balance performance and the significant predictors of balance performance from our MRA, i.e., S1-M1 and Lob VII-M1 connectivity (Table 5). Our results show that the relationship between connectivity and balance was stronger in the low cognition group, particularly for S1-M1 connectivity and balance (Figure 4).

**Figure 4:**
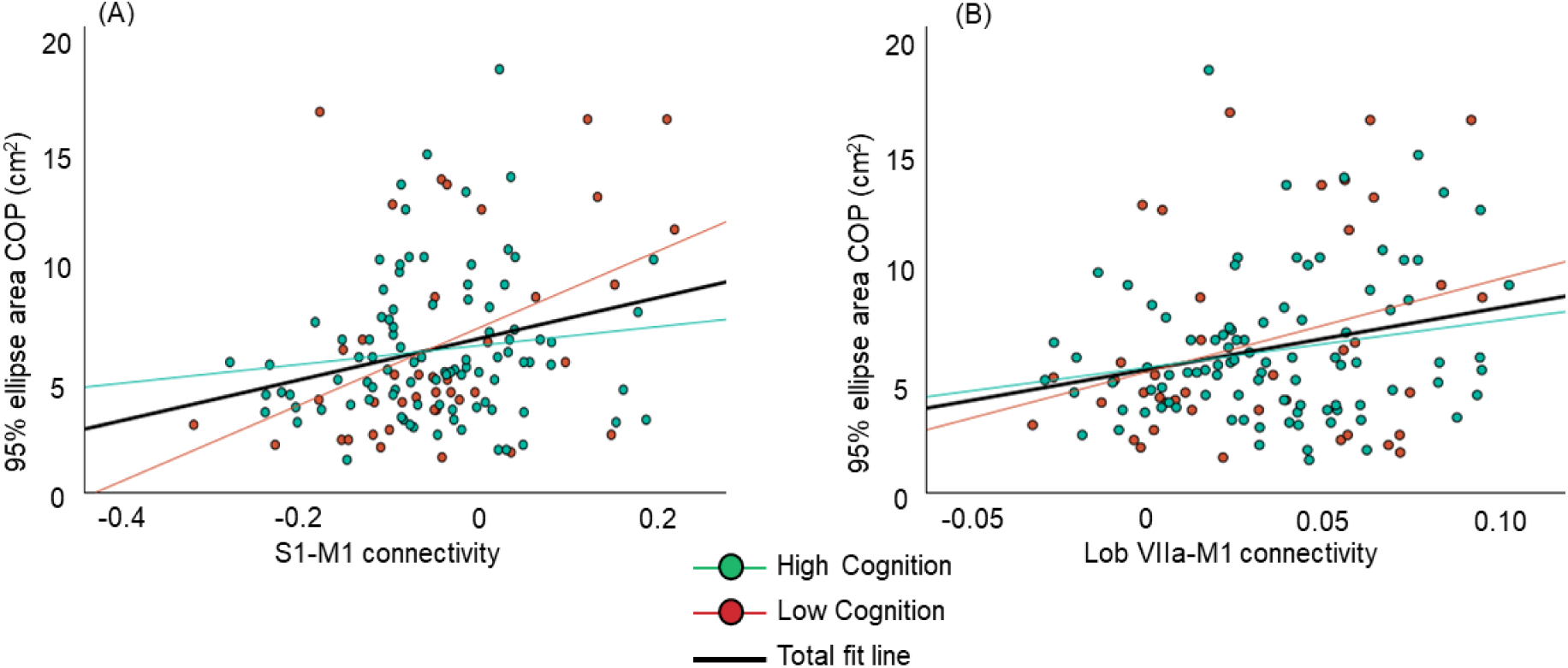
Bivariate correlations between (A) balance performance on ECCB and S1-M1 connectivity, (B) balance performance on ECCB and Lobule VIIa-M1 connectivity showing subgroups by MOCA score (green = high cognition: 26-30, red = low cognition: 20-25)

**Table 5:** Bivariate correlations between balance performance and connectivity split by subgroups based on cognition.

|  |  | S1-M1 | Lob VII-M1 |
| --- | --- | --- | --- |
| Low cognition group (MOCA scores between 20-25) | Pearson Correlation | 0.431 | 0.309 |
|  | P value | 0.006 | 0.053 |
| High cognition group (MOCA scores between 26-30) | Pearson Correlation | 0.127 | 0.201 |
|  | P value | 0.223 | 0.053 |

**Table 6:**
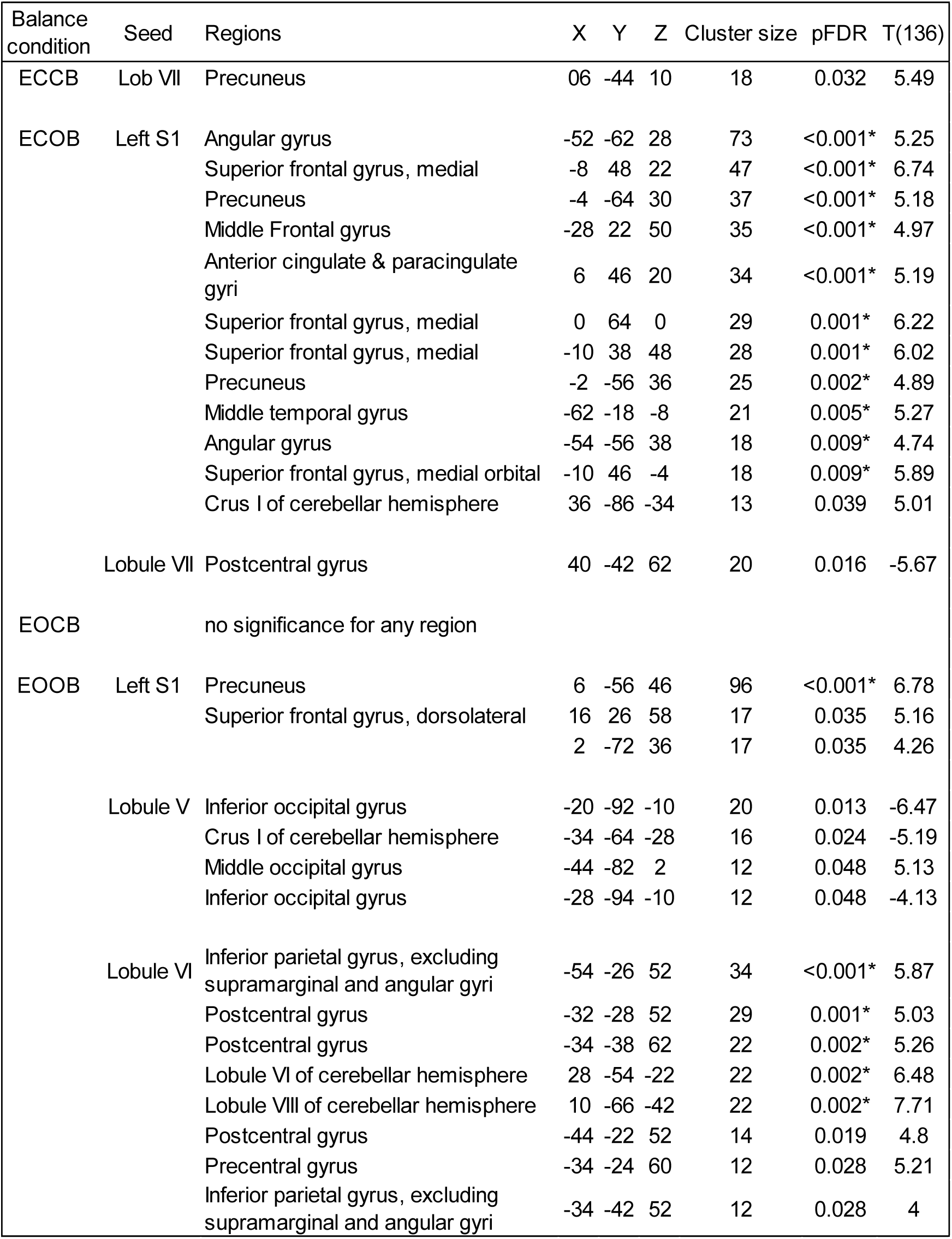
Coordinates of the brain regions showing significant relationship between performance on each of the four balance conditions and whole brain connectivity of S1 and cerebellum. Asterisks indicate FDR corrected p< 0.01.

We then performed moderation analyses using cognition (high and low cognition groups) as a moderator on the two significant correlations i.e., (1) performance on ECCB (dependent variable) with S1-M1 connectivity (independent variable) and (2) performance on ECCB (dependent variable) with Lobule VII-M1 connectivity (independent variable). Our moderation analysis showed that there was a significant interaction found by cognition on S1-M1 connectivity and balance (b= -0.0002, p =0.038), showing that cognition influences the relationship between connectivity and postural sway during ECCB. Further, age did not significantly directly moderate relationship between S1-M1 connectivity and balance (b= 0.0002, p= 0.4517) nor did it moderate the effect of cognition (moderated moderation) on the relationship between S1-M1 connectivity and balance (b= 0.0002, p = 0.2517). Cognition did not significantly moderate the relationship between Lobule VII-M1 connectivity and balance.

### 3.4. Exploratory analysis of the relationship between balance performance and the whole brain connectivity of S1 and cerebellum

Table XYZ shows the results of our exploratory analyses looking at the relationship between balance performance on all four conditions and the connections of S1 and cerebellum with the whole brain. Our results show that the connectivity of S1 with precuneus is significantly related to balance performance on the easiest condition (EOOB). On the more difficult ECOB condition, in addition to its connections with precuneus, connections of S1 with multiple other cortical regions, including angular gyrus, cingulate gyrus, superior frontal gyrus, middle temporal gyrus are significantly related to balance performance. For cerebellar connectivity, performance on the easier EOOB condition is significantly related to the connectivity of Lob VI with sensory cortex, including postcentral gyrus and inferior parietal gyrus. Further, balance performance on more difficult conditions ECOB and ECCB was related to the connectivity of Lobule VII with postcentral gyrus and precuneus, though this relationship did not meet statistical significance using the additional threshold of p_FDR_<0.01 for multiple comparisons.

## 4. Discussion

This study examined the relationship between connectivity of two brain regions that are critical to standing balance, namely, S1 and the cerebellum with M1. We found that S1-M1 and Lobule VIIa-M1 connectivity had significant correlations with performance on a standing balance task, indicating that connections from somatosensory cortex and cerebellum to motor cortex are related to balance control. Further, we also showed that cognition moderated the relationship between S1-M1 connectivity and standing balance, such that in individuals with worse cognition, the relationship between S1-M1 connectivity and balance was significantly stronger compared to those with higher cognition. This is one of the first studies to directly investigate the moderating effect of cognition when examining the relationship between brain connectivity and balance performance and adds to our understanding of the complex relationships between declines in network connectivity, balance, and cognition with aging. The implications of this work are discussed below.

### Greater S1-M1 and Lobule VII-M1 connectivity is related to poorer balance performance

Our results show that S1-M1 and Lobule VII-M1 connectivity was significantly related to balance performance on the ECCB condition. Higher S1-M1 and Lobule VII-M1 connectivity was correlated with higher postural sway, implying worse balance. Thus, our hypothesis is partially fulfilled in that connections from somatosensory cortex and cerebellum to motor cortex are important for standing balance, but that relationship is inverse i.e. greater connectivity is related to poorer balance performance. While the literature often discusses fcMRI differences in advanced age in the context of lower connectivity or decreases over time,(Andrews-Hanna et al., 2007; Onoda et al., 2012) including in our own work (Bernard et al., 2013; Ballard et al., 2022), heterogeneity is also seen in these connectivity patterns (Ferreira and Busatto, 2013; Roski et al., 2013; Farràs-Permanyer et al., 2019). That is, while decreased connectivity relative to young adults is the most widely reported pattern, areas of increased connectivity are also seen, including in our own work (Bernard et al., 2013, 2021). Notably for our work here, this has been demonstrated in motor circuits and networks in a large sample (n=191), including in cerebellar connectivity (Seidler et al., 2015b). Further, there were associations with motor tapping performance that linked higher connectivity with higher reaction time (slower performance) (Bernard et al., 2013), which in many ways parallels our findings here with respect to postural control. A potential explanation for the increase in functional connectivity that accompanied poorer balance performance, particularly with aging, is that this is indicative of an increase in functional recruitment of additional brain networks to involve additional neural resources to maintain upright balance, despite a decay in the brain networks more generally, as a means of compensation (Cabeza et al., 2019). Of course, this balance assessment was conducted offline in a separate session, but the resting state signal may reflect this compensatory attempt. Prior neuroimaging work lends support to this concept of compensation, wherein well-preserved structural connectivity was accompanied by low connectivity between networks while greater structural declines with aging were accompanied by a functionally maximally connected nervous system (Stumme et al., 2022).

### Cognition moderated the relationship between S1-M1 connectivity and balance

Additionally, we found that the strength of relationship between S1-M1 connectivity and balance differed depending on cognition and was significant in individuals with low levels of cognition but not in individuals with higher levels of cognition. Advanced age leads to an overall decline in fluid cognition (Park et al., 2001; Park and Reuter-Lorenz, 2008) and balance (Horak et al., 1989). Further, cognitive impairment is associated with worse balance performance (Shin et al., 2011). Our results add to the current evidence on the relationship between cognition and balance by showing that for the same level of connectivity, an individual with lower cognition had greater postural sway compared to an individual with higher cognition. It may be that individuals with poor cognition need increased recruitment of brain regions (compensation for cognitive declines) and in turn, higher wiring costs, which would be associated with increased functional connectivity. This is supported by the compensation related utilization of neural circuits hypothesis that purports that networks in an aging brain need to “work harder” to make up for processing declines in the same network or elsewhere in the brain (Reuter-Lorenz and Cappell, 2008). Further, the balance task used in our analyses i.e., eyes closed with closed base of support, was a challenging task that required individuals to rely more on their sensory input and adapt their body sway in response to these sensory cues. It is possible that given the difficulty of this task, individuals with better cognition were better able to use the neural resources available to them to adapt to the demands of this task. This is in line with prior work that has shown that there is age-related extra-recruitment of neural resources under taxing conditions, such as when novel, weak, and complex rules must be acquired (Vallesi et al., 2011). Here we speculate that this may extend to the motor domain, in this case in a more challenging balance task.

### Cognition did not moderate the relationship between cerebellar-M1 connectivity and balance

In addition to its role in motor control, the cerebellum, particularly Lobule VII is known to play a role in cognition as well (Stoodley, 2012a; Schmahmann, 2019). Hence, it was surprising that cognition did not moderate the relationship between Lobule VII-M1 and balance performance. One explanation, especially given the dominant role Lobule VII plays in cognition, is that the Lobule VII-M1 connectivity itself captured the cognitive contributions in our moderation model, leaving little left for an additional moderator (such as MOCA score) to explain. An alternative explanation is that it is possible that cognition moderates the relationship between balance performance and cortico-cortical networks (such as S1-M1 connectivity noted above) but not cerebello-cortical networks. In particular, the manner in which the cerebellum contributes to cognitive function may be different from cortico-cortical circuits. While cerebellar contributions to motor control are thought of in context of an internal model, a similar framework has been hypothesized regarding cerebellar contributions to non-motor functions such as cognition as well (Ito, 2008; Ramnani, 2014). Like how cerebellum acts on efference copies of motor commands that originate in M1, the cerebellum is also hypothesized to act on efferent copies of “thoughts” that might originate in the cerebral cortex. Thus, the nature of cerebellar contributions, which is to refine and adapt the cognitive output, is different from contributions of the cortico-cortical circuits, which are more directly involved in generating cognitive output. While this difference in cortico-cortical circuits and cerebello-cortical circuits may not be entirely responsible for the lack of moderation seen in our results, it may offer some insight into the difference in the effect of cognition on cortico-cortical versus cerebello-cortical circuits.

### Age did not influence the relationship between connectivity and balance

We had originally hypothesized that lower connectivity would be related to worse balance based on the age differences in both functional connectivity and balance performance. However, our data showed that not only was age not a key factor in influencing the relationship between connectivity and balance performance but also age did not affect how cognition moderated the relationship between connectivity and balance performance. Thus, cognition rather than age may be a critical factor that influences the balance performance in middle aged and older adults. Also, given that our moderation results were from the entire sample, ranging in age from 35 to 86, and not just older adults, our findings highlight the importance of the effect of cognitive loss on balance control not only in older adults but also in middle-aged adults. Thus, cognitive decline may start influencing balance control as early as the middle age (35 years onwards), even though the effects may be more apparent in the older adult group.

### Whole brain analysis of S1 and cerebellar connectivity

Exploratory whole brain analyses showed that while the balance performance on EOOB is related to connectivity of S1 with precuneus, a region known to play an important role in complex cognitive functions as well as sensorimotor integration and visuo-spatial processing (Tanglay et al., 2022; Dadario and Sughrue, 2023), with increasing balance complexity, such as on ECOB, balance performance is related to S1 connectivity with additional cortical regions, such as angular gyrus, cingulate, frontal and temporal lobe, which are all involved in several executive and higher order cognitive functions (Devinsky et al., 1995; Boisgueheneuc et al., 2006; Seghier, 2013; Xu et al., 2019). Similarly, with respect to cerebellar connectivity, connectivity of the lobules primarily involved with motor behavior, such as Lobule V and Lobule VI, is related to performance on the easier EOOB condition (Habas et al., 2009). With more difficult balance conditions, such as with ECOB and ECCB, there is a shift to Lobule VII, which is known to play a role in cognitive function (Stoodley, 2012b). These results indicate that as the balance conditions get more demanding, there may be a need for greater cognitive resources to maintain upright balance and thus, support our moderation analyses by showing that poorer cognition may influence balance performance.

It is also interesting to note that our results show that the connectivity between S1 and cerebellum itself, specifically, Lobules VI and VII, is related to balance performance. CB has been shown to modulate sensorimotor behavior through Purkinje cell projections to several cortical regions, including the frontal and parietal lobe via the thalamus (Strick et al., 2009; Lindeman et al., 2020). Our findings lend support to these connections of the cerebellum with non-motor regions, particularly the sensory cortex, and underscore the contribution of cerebellum to balance control.

### Clinical implications

Altogether, our results show that the networks between S1-M1 and cerebellum-M1 are critical for upright balance. While the role of both S1 and cerebellum in upright balance control has been investigated previously, our work highlights the contributions of network connectivity of these two regions. This is particularly important when considering the effects of interventions such as non-invasive brain stimulation to these regions, wherein the observed changes in behavior may not be solely due to stimulation of S1 or cerebellum but also due to changes in the interactions of these regions with other brain regions.

Our work also suggests that cognition, network connectivity, and balance have a complex relationship. From a balance rehabilitation perspective, screening for cognitive declines and using a comprehensive multi-disciplinary approach that addresses all these factors is needed to impact balance control and reduce fall risk. Our results underscore the need for interventions that not only focus on improving sensory and motor systems for balance rehabilitation but also address cognitive declines that accompany aging. Furthermore, given that our results highlight the role of network connectivity of S1 and cerebellum in maintaining upright balance control, future work can investigate the utility of S1 and cerebellum as potential targets for interventions such as non-invasive brain stimulation.

## 5. Limitations

While this work provides valuable insight into the complex relationship between sensorimotor and cerebellar-M1 connectivity, it is not without limitations. We determined our low and high cognition groups based on MOCA scores. Although the MOCA is a commonly used screening tool that has been shown to be a valid and reliable measure of cognitive function, it is not as comprehensive as the full battery of neuropsychological tests used to diagnose individuals with mild cognitive impairment (MCI). Hence, our conclusions can only be applied to individuals with low scores on the MOCA and cannot be generalized to individuals with a diagnosis of MCI. However, while the above may be a limitation, it may also aid in the application of this knowledge to clinical practice and in rehabilitation, as the MOCA is a clinically feasible tool that is easy to administer for a health care provider or a physical therapist whereas obtaining a clinical diagnosis of MCI can take time. Also, our data are cross-sectional and correlational, and hence, we cannot make any causal claims. Further investigations tracking functional connectivity, cognition and balance performance across time would be helpful in providing a greater insight into the complex triad of connectivity, cognition, and balance.

In conclusion, our results suggest that connections of the primary somatosensory cortex and cerebellum with primary motor cortex play an important role in balance control. Cognition moderated the relationship between sensorimotor connectivity and balance, such that individuals with lower cognitive function show a stronger relationship between sensorimotor connectivity and balance. Age did not moderate the relationship between connectivity and balance, nor did it indirectly influence the effect of cognition on this relationship. Our results are an important first step in identifying primary sensory cortex and the specific cerebellar lobules as targets for neuromodulation to improve balance performance.

## 6. Acknowledgements

This work was supported by the National Institute on Aging R01AG064010 to JAB. This work was further supported in part by the Texas Virtual Data Library (ViDaL), funded by the Texas A&M University Research Development fund.

