## Supplementary Table 1 and Supplementary Table 2 for "Relationships between balance performance and connectivity of motor cortex with primary somatosensory cortex and cerebellum in middle aged and older adults"

Supplementary Table 1: Pearson’s correlation coefficient and p values for the relationship between balance (three balance conditions: EOOB, ECOB, and EOCB) and connectivity (S1-M1, Lobule V-M1, Lobule VI-M1, Lobule VII-M1, Lobule VIII-M1 and Lobule X-M1.

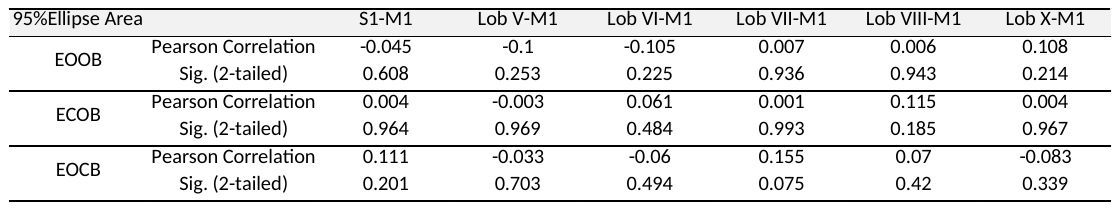

Supplementary Table 2: Pairwise Comparisons between the 95% COP ellipse area for the four balance conditions

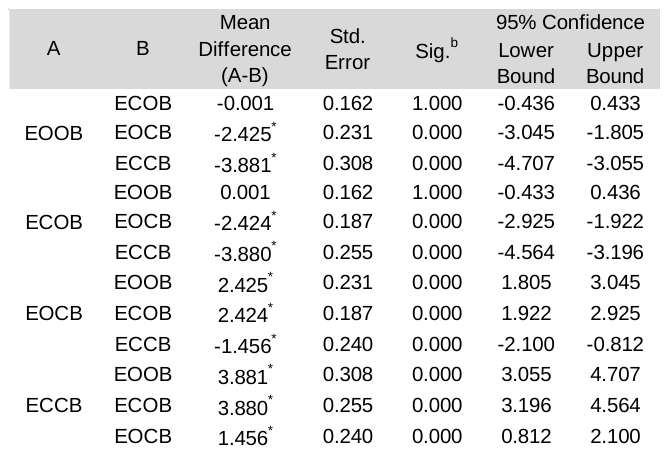

Based on estimated marginal means

*. The mean difference is significant at the .05 level.

b. Adjustment for multiple comparisons: Bonferroni.
